# Subtle Facial Skin Color Changes Enhance the Human Likeness of Artificial Agents: Insights from Pulse-Wave-Derived Modulation

**DOI:** 10.64898/2026.05.22.726810

**Authors:** Saya Nikaido, Tomoko Isomura

## Abstract

Recent studies have shown that implementing explicit social cues, such as gaze, facial expressions, and gestures, in artificial agents can improve impressions of these agents. However, humans may also use implicit physiological cues, such as facial coloration and cardiac information, in social perception. The present study examined whether subtle facial skin color changes derived from pulse-wave signals enhance the perceived human likeness of artificial agents and whether this effect depends on the type of agent, the temporal characteristics of the signal, or individual differences in interoceptive sensibility. Participants observed morphed face stimuli created from artificial agents and human faces and judged whether each stimulus appeared human-like or robot-like. In Experiment 1, skin color changes based on human-derived pulse-wave signals enhanced perceived human likeness for a highly human-like agent, but not for a less human-like agent. In Experiment 2, perceived human likeness was enhanced not only by pulse-wave-derived facial skin color changes but also by sinusoidal color changes matched to the pulse-wave signals in mean amplitude and number of peaks. Interoceptive sensibility did not significantly moderate the effect of facial skin color changes on perceived human likeness. These findings suggest that periodic facial skin color changes with physiologically plausible temporal characteristics, rather than temporal dynamics specific to pulse-wave signals, may serve as cues that enhance the perceived human likeness of artificial agents.

## Introduction

Research in human–agent interaction (HAI) has shown that people often respond to artificial agents, such as robots and virtual avatars, as if they were social beings (Nass et al., 1994; Nass & Moon, 2000). Moreover, observers often adopt an intentional stance toward artificial agents, interpreting their behavior in terms of underlying beliefs, desires, and intentions (Dennett, 1989). As the attribution of such internal states to artificial agents is central to successful human–agent interaction (Schellen & Wykowska, 2019), researchers have sought to identify and implement cues that promote such attributions. Accordingly, a growing body of research has focused on social cues such as facial expressions, gaze, and body language that facilitate observers’ attribution of internal states to artificial agents (Breazeal, 2003; Admoni & Scassellati, 2017; Beck et al., 2012; Feine et al., 2019).

Although most cues studied in HAI research to date are explicit social cues, recent psychological research suggests that humans also use subtle physiological cues to infer others’ internal states. For example, Benitez-Quiroz et al. (2018) demonstrated that distinct facial color patterns are associated with different emotional states and that facial coloration can provide information about others’ emotions, independently of morphological cues such as facial expressions. Furthermore, Galvez-Pol et al. (2022) and Arslanova et al. (2022) demonstrated that observers can match heartbeat rhythms to the corresponding individual by viewing faces. The authors suggested that observers may use, at least in part, transient facial color changes synchronized with heartbeats. Together, these findings suggest that physiological information reflected in subtle changes in facial skin color can be conveyed to observers and may serve as a cue for inferring others’ internal states.

These findings raise the possibility that heartbeat-related physiological cues may also influence the perception of artificial agents. Indeed, McDuff and Nowara (2021) investigated the effects of pulse-wave-derived facial skin color changes on impressions of synthetic avatars. They applied skin color changes based on pulse-wave signals to the faces of synthetic avatars created from human images. Although the color changes were imperceptible, impressions of naturalness and aliveness were enhanced relative to a no-pulse condition. Furthermore, ratings of naturalness, lifelikeness, consciousness, and aliveness were higher in the pulse-wave condition than in a sinusoidal condition in which the facial color changed toward red in a sinusoidal rhythm. This suggests that impressions of artificial agents may be influenced not merely by periodic changes in facial coloration, but by biologically plausible temporal dynamics embedded in these changes. McDuff and Nowara (2021) also reported that increasing the avatar’s simulated heart rate increased perceptions of the avatar’s arousal, suggesting that cardiac signals can influence the attribution of arousal-related internal states to artificial agents.

Although this pioneering study provides important insights into the effects of pulse-wave-derived facial skin color changes on impressions of artificial agents, several important questions remain. First, the stimuli used in that study were highly human-like avatars, and it remains unclear whether similar effects would be observed in agents with lower levels of human likeness. Second, because the study primarily relied on comparative judgments between skin-color conditions, it remains unclear how pulse-wave-derived skin color changes influence perceptions of the human likeness of individual agents. Third, the pulse-wave and sinusoidal conditions differed not only in their temporal dynamics but also in the naturalness of the skin color changes. Consequently, it remains unclear how sinusoidal skin-color changes would influence impressions of agents when implemented in a more naturalistic manner. Finally, observer-related factors that may influence sensitivity to these effects have yet to be examined.

Building on the findings of McDuff and Nowara (2021), the present study examined the effects of pulse-wave-derived facial skin color changes on perceived human likeness in artificial agents. Our first aim was to identify the conditions under which this effect emerges. Specifically, we examined whether this effect generalizes across agents with different levels of human likeness, whether it is specific to biologically derived pulse-wave dynamics or can also emerge from artificially generated sinusoidal dynamics when implemented in a naturalistic manner.

Our second aim was to investigate observer-related factors associated with sensitivity to these effects. In particular, we focused on interoceptive sensibility and examined whether individuals with greater awareness of bodily signals are more likely to perceive artificial agents with pulse-wave-derived skin color changes as human-like. Previous research has suggested that interoceptive sensibility is associated with aspects of social perception, including the inference of others’ emotional states and empathy (Hübner et al., 2021; Canino et al., 2024). This association may arise because individuals with greater awareness of their own bodily signals are more likely to incorporate physiological information into social judgment. From this perspective, physiological cues implemented in artificial agents may have a greater influence on social perception among observers with higher interoceptive sensibility, potentially promoting the attribution of internal states and enhancing the perceived human likeness of the agent.

To address these aims, we employed a psychophysical paradigm using face stimuli morphed along a continuum between human and artificial agent identities. The stimuli consisted of 8-s videos in which subtle facial skin color changes were implemented or omitted. In online experiments, participants judged whether each stimulus appeared more human-like or robot-like. This approach enabled a fine-grained assessment of how pulse-wave-derived skin color changes influence perceived human likeness across different levels of agent human likeness. In Experiment 1, we examined whether the effect of pulse-wave-derived facial skin color changes varied across agents with different levels of human likeness. In Experiment 2, we examined whether this effect was specific to human-derived pulse-wave signals by comparing them with artificially generated sinusoidal signals. In both experiments, we assessed participants’ interoceptive sensibility using a questionnaire. We hypothesized that artificial agents would be perceived as more human-like when pulse-wave-derived facial skin color changes were present than when they were absent. We also explored whether the magnitude of this effect would vary depending on the agent’s level of human likeness and the temporal dynamics of the implemented skin color changes. We further hypothesized that this effect would be stronger among participants with higher interoceptive sensibility.

### Experiment 1

The aim of Experiment 1 was to examine whether pulse-wave-derived facial skin color changes enhance the perceived human likeness of artificial agents, and whether this effect varies across artificial agents with different levels of human likeness. We used two artificial agents, ERICA and CommU, which differ in their degree of human likeness. Participants viewed morphed human-agent faces with or without pulse-wave-derived skin color changes and judged whether each stimulus appeared more human-like or robot-like.

## Methods

### Participants

Participants were recruited through CrowdWorks and assigned to either the CommU condition or the ERICA condition. A total of 620 participants completed the task. Twenty-two participants were excluded from the analysis because their questionnaire responses indicated that they had not followed the task instructions. In addition, to identify participants with atypical response patterns across morph levels, Pearson’s correlation coefficient between morph level and human likeness judgment was calculated for each participant. The correlation coefficients were then Fisher *z*-transformed, and participants whose transformed coefficients fell outside the mean ± 3 SD of the distribution were excluded. Six participants met this criterion. The final sample comprised 592 participants: 302 in the CommU condition (121 men, 175 women, and 6 who selected “other” or “prefer not to answer”; mean age = 41.89 years, SD = 10.50) and 290 in the ERICA condition (111 men, 174 women, and 5 who selected “other” or “prefer not to answer”; mean age = 41.24 years, SD = 10.98).

### Stimuli

The stimuli consisted of 8-s videos of faces morphed between human and artificial agent identities, presented with and without pulse-wave-derived facial skin color changes. Two artificial agents were used: ERICA, a geminoid with a highly human-like appearance, and CommU, which has a relatively robot-like appearance. Both agents were developed as part of the ERATO Ishiguro Symbiotic Human-Robot Interaction Project.

Three human faces were selected from the Yonsei Face Database (Chung et al., 2019) based on a preliminary experiment to ensure that changes in perceived human likeness across morph levels were comparable across human identities. Because ERICA has a female appearance, only female faces were used to avoid introducing sex-related morphological differences (e.g., facial hair) into the morphed stimuli.

Seven morph levels were defined for each human-agent face pair. These levels were determined based on a second preliminary experiment such that levels 1, 4, and 7 corresponded approximately to 0%, 50%, and 100% probabilities of being judged as human-like, respectively. Intermediate levels were logarithmically spaced between these anchor points. The resulting human-face proportions for morph levels 1–7 were 60%, 73%, 77%, 80%, 83%, 87%, and 100% for CommU, and 0%, 36%, 46%, 50%, 54%, 64%, and 100% for ERICA. Skin tone was adjusted using Adobe Photoshop so that the hue remained constant across all morph levels for each human face.

Pulse-wave signals were recorded photoplethysmographically from the right cheek of one healthy adult during an 8-s recording following a 3-min resting period. Signals were sampled at 1000 Hz using a pulse wave sensor [AP-C030(A)] connected to a POLYMATE AP5148 system (Miyuki Giken Co., Ltd.).

Pulse-wave-derived skin color changes were implemented by modulating the RGB values of each stimulus according to the recorded pulse-wave signal, and 8-s videos were generated at 30 fps. The RGB modulation coefficients were set to 0.23, 0.41, and 0.36 for the red, green, and blue channels, respectively, following McDuff and Nowara (2021). These coefficients reflect the relative signal strength in each color channel when pulse-related optical changes in the skin are captured by an RGB camera (Lin, 2014) (Figure 1). The implemented skin color changes were designed to be sufficiently subtle and were not readily noticeable under normal viewing conditions. Control stimuli were identical except that no pulse-wave-derived skin color changes were applied.

**Figure 1.**
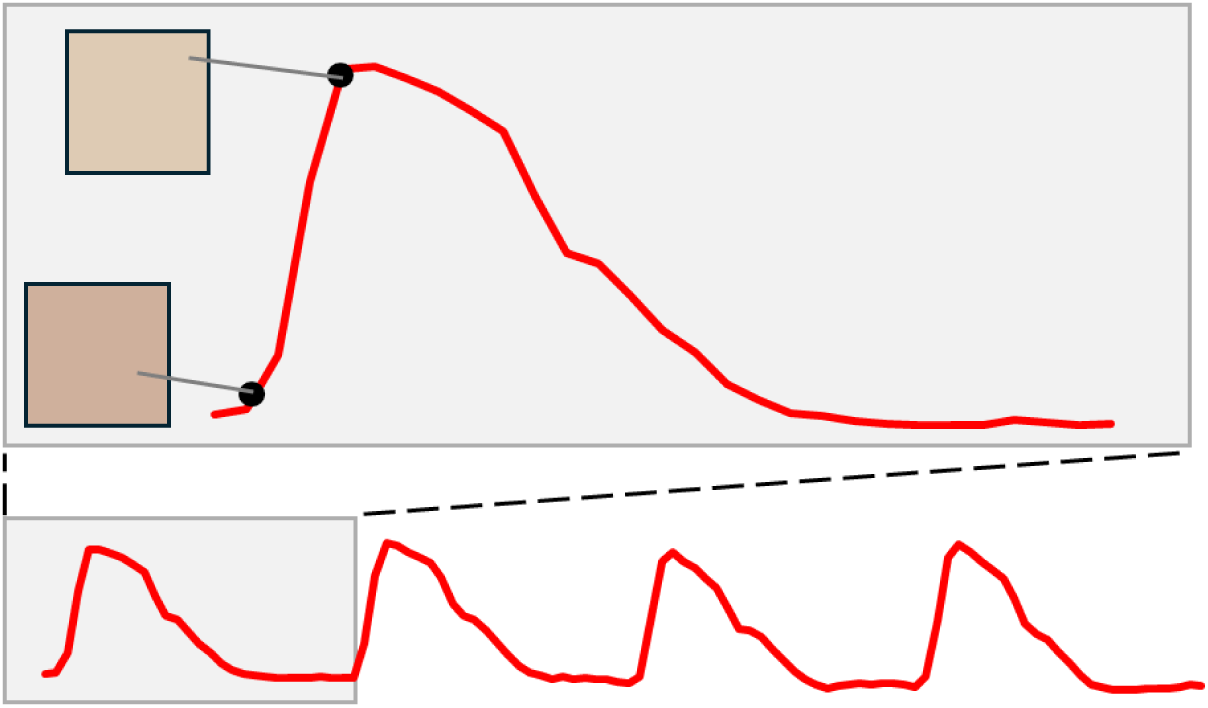
Schematic illustration of the implementation of pulse-wave-derived facial skin color changes. Facial skin color varied in synchrony with the recorded pulse-wave signal. The magnitude of the color changes is exaggerated in this figure for illustrative purposes.

### Procedure

#### Human-likeness judgment task

Participants completed an online human-likeness judgment task implemented in Qualtrics. Each participant was assigned to one artificial agent (CommU or ERICA) and one of three human-face identities. The use of multiple human-face identities was intended to reduce the possibility that the results reflected idiosyncratic responses to a particular face. Within the assigned agent-face pair, participants viewed stimuli from both color-change conditions (pulse-wave vs. no-pulse). In each block, participants observed 14 video stimuli, each lasting 8 s, and evaluated whether the target appeared “human-like” or “robot-like” using a two-alternative forced-choice task. The experiment consisted of four blocks, with stimulus order randomized within each block. To reduce variability in viewing conditions, participation was restricted to mobile devices such as smartphones and tablets. Participants were instructed to complete the experiment indoors, minimize screen glare, and set their screen brightness to the maximum level.

#### Questionnaires

After completing the human-likeness judgment task, participants completed all 32 items of the Japanese version of the Multidimensional Assessment of Interoceptive Awareness (MAIA-J; Mehling et al., 2012; Shoji et al., 2014) to assess interoceptive sensibility. Although this version consists of eight subscales, subscale scores were calculated according to the 25-item, six-factor structure of the MAIA-J validated by Shoji et al. (2018). Participants then answered two questions about their experience during the experiment. First, they provided a free-text response describing which aspects of the stimuli they focused on when making human-likeness judgments. Second, they indicated whether they had noticed any changes in facial skin color by selecting one of three response options: “noticed,” “did not notice,” or “other”. In addition, participants were given an optional free-text field to describe anything they had noticed regarding changes in facial skin color. The questionnaires were created and administered using Qualtrics.

### Data Analysis

#### Testing the Effect of Pulse-Wave-Derived Facial Skin Color Changes

To examine whether pulse-wave-derived facial skin color changes enhanced perceived human likeness, we fitted generalized linear mixed models (GLMMs) with a binomial distribution and logit link function. Trial-level binary judgments, coded as human (1) or robot (0), were used as the outcome variable. Separate models were fitted for each artificial agent. Trial number and morph level were centered before analysis.

First, a baseline model was fitted including age, sex, trial number, morph level, and human face identity as fixed effects. The model also included participant-specific random intercepts and random slopes for trial number and morph level, as well as a random intercept for stimulus. The baseline model was specified as follows:

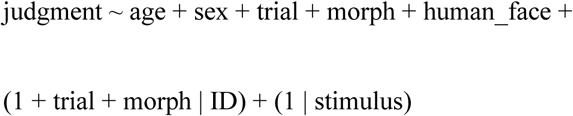

Next, to examine the effect of pulse-wave-derived skin color changes, a main-effects model was fitted by adding color-change condition as a fixed effect to the baseline model. The main-effects model was specified as follows:

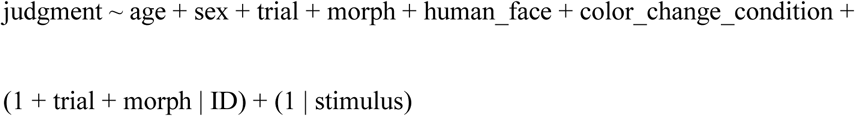

A likelihood ratio test (LRT) was used to assess whether adding color-change condition significantly improved model fit.

Furthermore, to examine whether the effect of pulse-wave-derived facial skin color changes was consistent across the human faces used for morphing, an interaction model was fitted by adding the interaction between color-change condition and human face identity to the main-effects model.

#### Examining the moderating effect of interoceptive sensibility

We next examined whether interoceptive sensibility moderated the effect of pulse-wave-derived facial skin color changes on human-likeness judgments. Based on the hypothesis that greater awareness of bodily signals and a stronger tendency to interpret them as meaningful information would be associated with greater use of physiological information into social judgment, we focused on four of the six MAIA subscales that were theoretically relevant to this hypothesis: Body Listening, Noticing, Attention Regulation, and Emotional Awareness. Body Listening reflects the tendency to actively listen to the body for insight; Noticing reflects awareness of bodily sensations; Attention Regulation reflects the ability to sustain and control attention to bodily sensations; and Emotional Awareness reflects awareness of the connection between bodily sensations and emotional states. Each subscale score was calculated as the mean of its constituent items. Prior to analysis, scores on these four subscales were centered and entered into the model as interaction terms with color-change condition.

The fitted model (hereafter referred to as the MAIA model) was specified as follows:

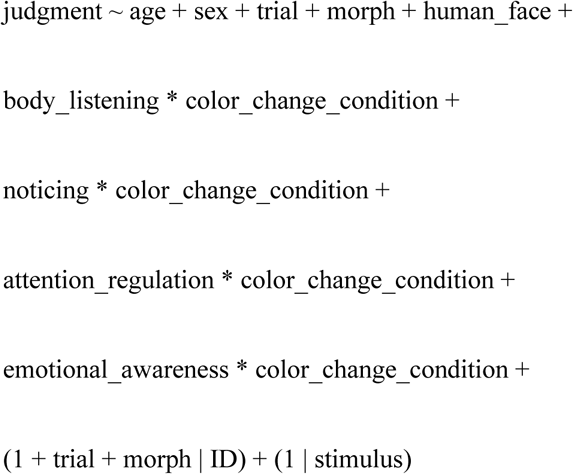

#### Examining the moderating effect of awareness of facial skin color changes

Although the skin-color changes were designed to be subtle and not readily noticeable, some participants reported noticing changes in facial skin color in their free-text responses. We therefore conducted an exploratory analysis to examine whether explicit awareness of these changes moderated the effect of color-change condition on human-likeness judgments. To test this, we fitted a model including the interaction between awareness and color-change condition (hereafter referred to as the color-change awareness model):

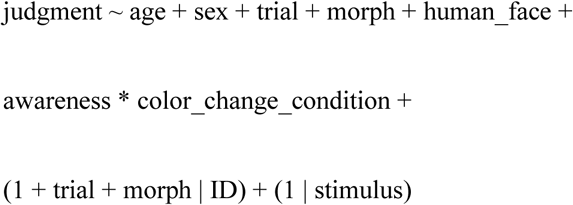

All GLMMs were fitted using the lme4 package in R (Bates et al., 2015).

## Results & Discussion

### Testing the Effect of Pulse-Wave-Derived Facial Skin Color Changes

In the CommU condition, adding color-change condition to the baseline model did not significantly improve model fit (χ²(1) = 0.17, *p* = .68). Morph level was positively associated with human-likeness judgments (*β* = 3.37, *z* = 19.04, *p* < .001), indicating, as expected, that stimuli were more likely to be judged as human as the proportion of the human face increased. However, the effect of color-change condition was not significant, indicating that pulse-wave-derived facial skin color changes did not reliably alter human-likeness judgments for CommU. Detailed results are presented in Table 1.

**Table 1.**
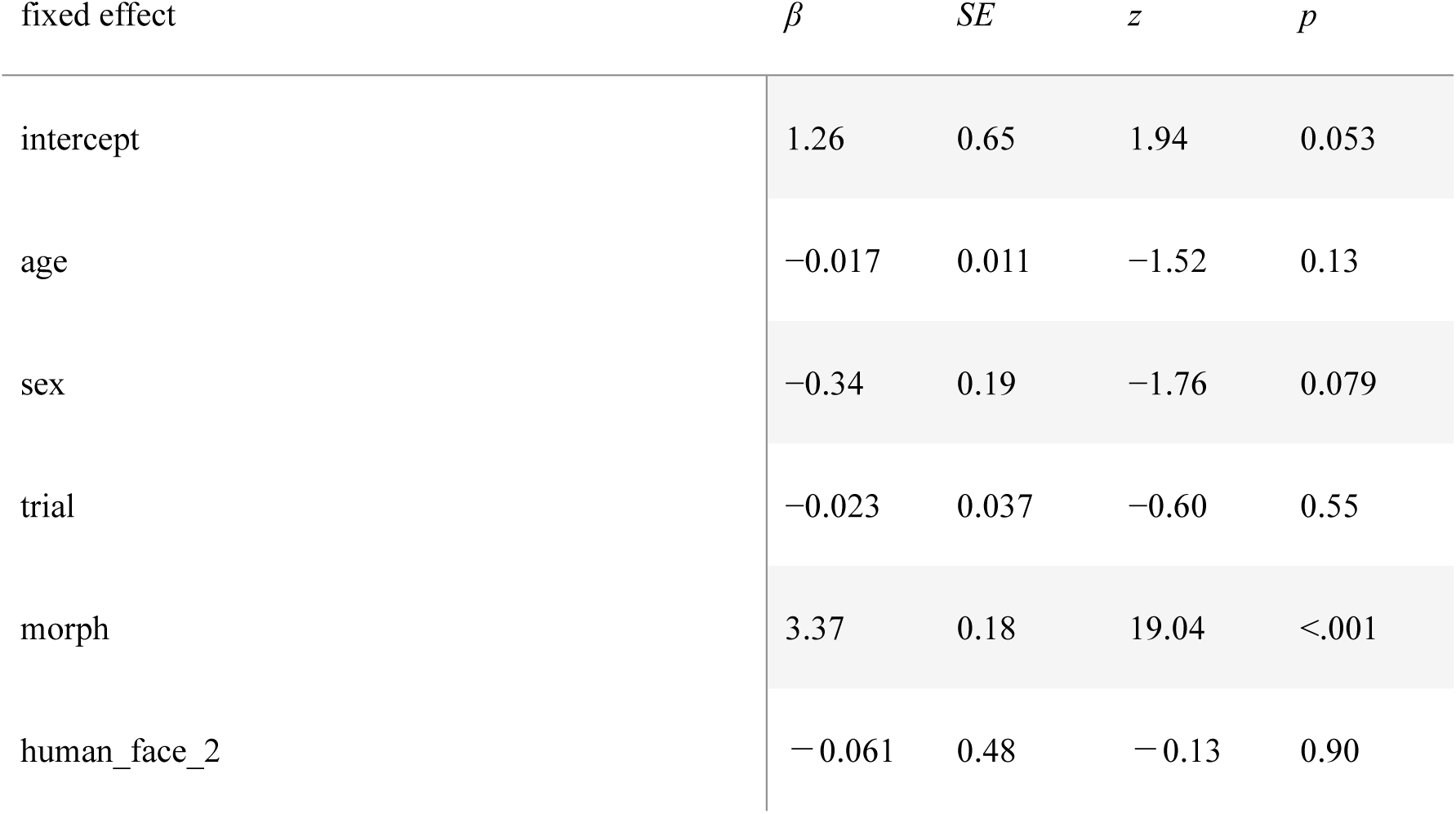
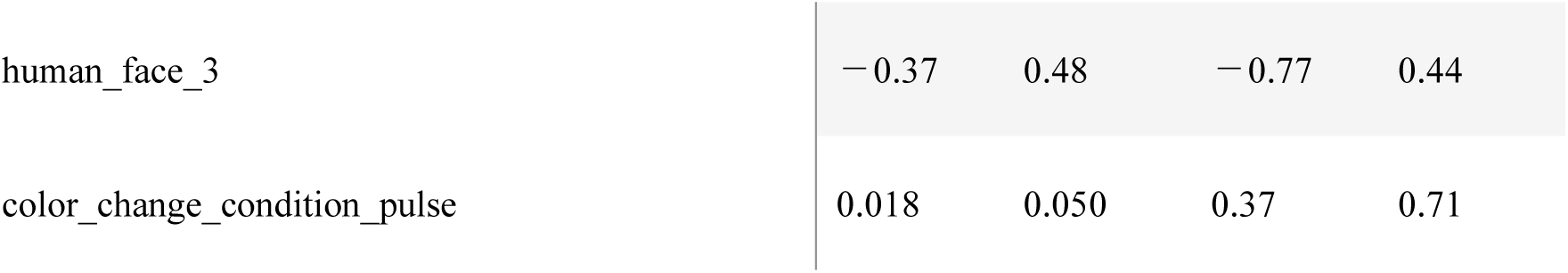
CommU condition main-effects model.

In the ERICA condition, adding color-change condition significantly improved model fit (χ²(1) = 13.21, *p* < .001). In addition to a significant effect of morph level (*β* = 3.12, *z* = 13.61, *p* < .001), color-change condition also had a significant effect (*β* = 0.19, *z* = 3.71, *p* < .001). Stimuli with pulse-wave-derived skin color changes were more likely to be judged as human than those without such changes. The odds of being judged as human were approximately 1.21 times higher in the pulse-wave condition than in the no-pulse condition (OR = 1.21, 95% CI [1.09, 1.34]). Detailed results are presented in Table 2.

**Table 2.** ERICA condition main-effects model.

| fixed effect | $\beta$ | $SE$ | $z$ | $p$ |
| --- | --- | --- | --- | --- |
| intercept | 1.50 | 0.74 | 2.04 | 0.042 |
| age | −0.002 | 0.010 | −0.18 | 0.86 |
| sex | −0.24 | 0.21 | −1.15 | 0.25 |
| trial | 0.061 | 0.037 | 1.65 | 0.099 |
| morph | 3.12 | 0.23 | 13.61 | <.001 |
| human_face_2 | 0.012 | 0.62 | 0.019 | 0.98 |
| human_face_3 | −0.28 | 0.61 | −0.46 | 0.65 |
| color_change_condition_pulse | 0.19 | 0.051 | 3.71 | <.001 |

Adding the interaction between color-change condition and human face identity to the model did not significantly improve model fit compared with the main-effects model (χ²(2) = 4.10, *p* = .13), suggesting the effect of skin color change was relatively consistent across the different human face identities used to construct the morph continua.

### Examining the moderating effect of interoceptive sensibility

We then tested whether individual differences in interoceptive sensibility moderated the effect of skin-color changes on human-likeness judgments. Four MAIA-J subscales were included in the model, together with their interactions with color-change condition. However, none of these interactions reached statistical significance for either CommU or ERICA. Thus, contrary to our hypothesis, interoceptive sensibility did not significantly moderate the effect of skin color changes on human-likeness judgments. Detailed results are presented in Tables 3 and 4.

**Table 3.**
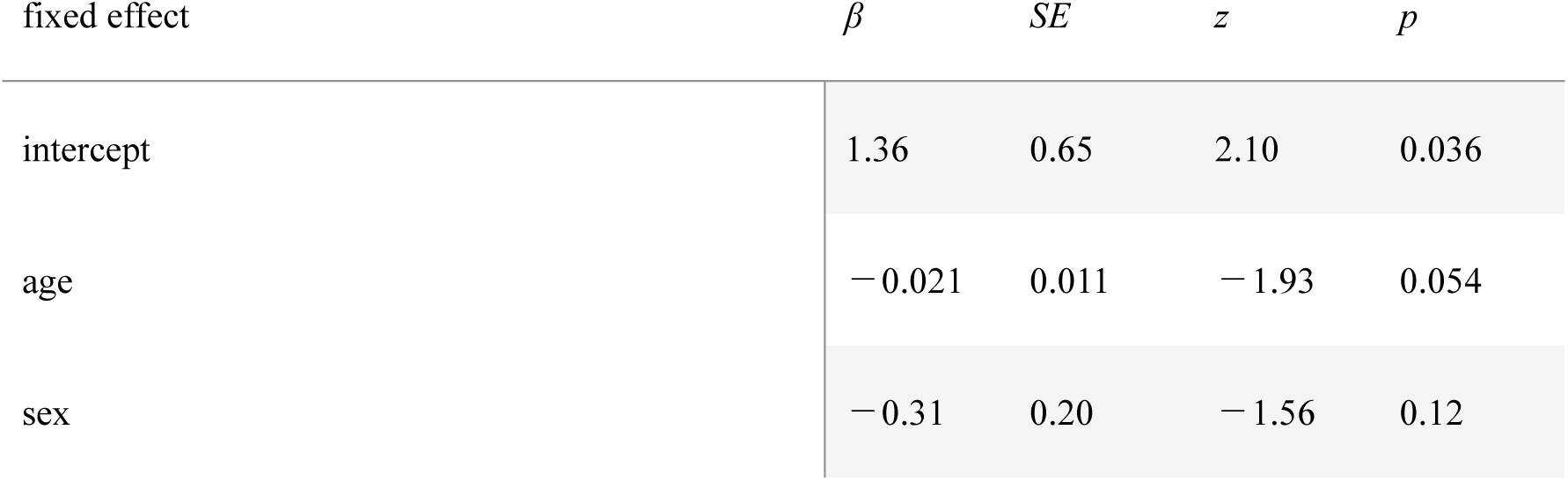
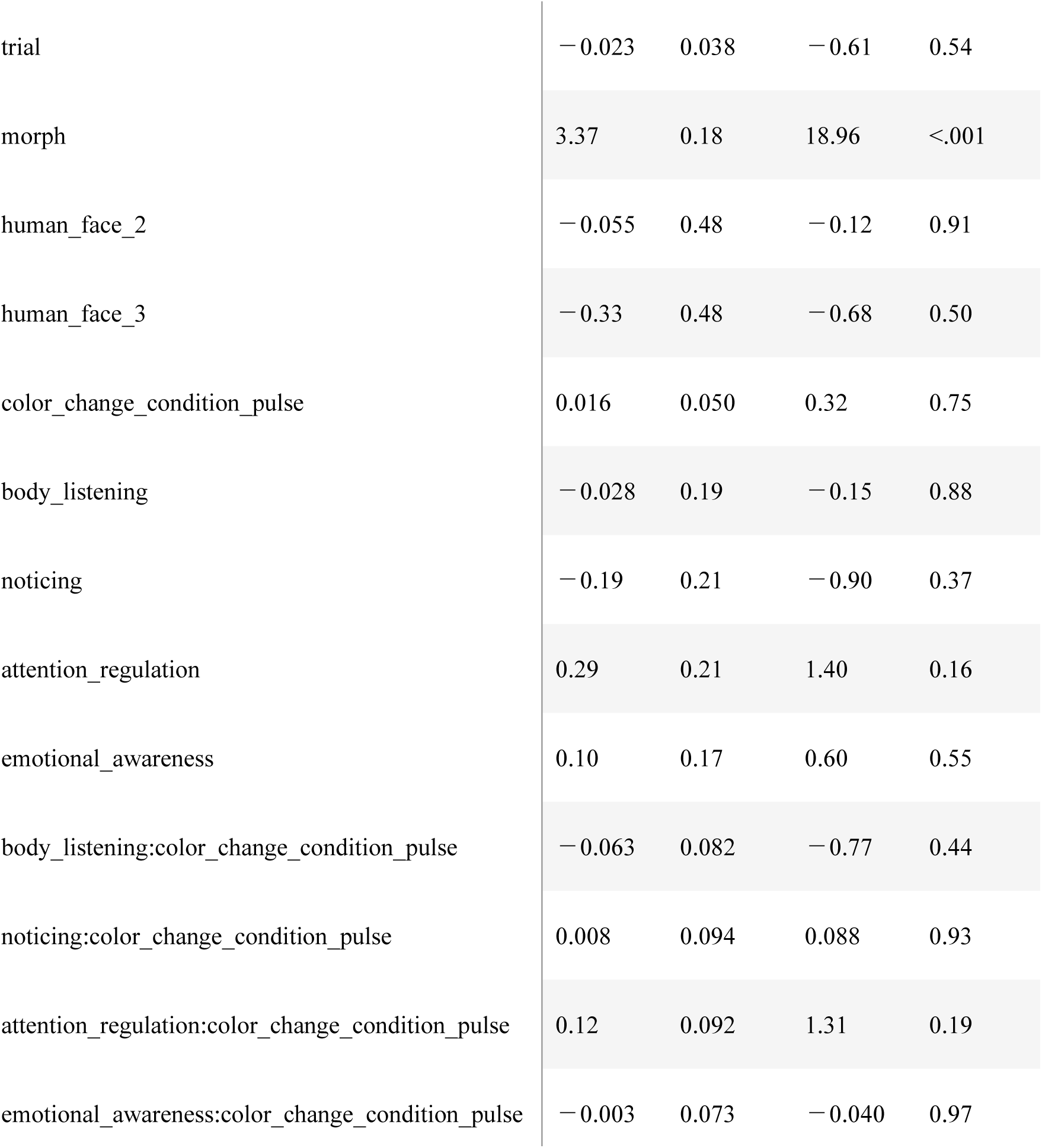
CommU condition MAIA model.

| fixed effect | $\beta$ | *SE* | *z* | *p* |
| --- | --- | --- | --- | --- |
| intercept | 1.36 | 0.65 | 2.10 | 0.036 |
| age | −0.021 | 0.011 | −1.93 | 0.054 |
| sex | −0.31 | 0.20 | −1.56 | 0.12 |
| trial | −0.023 | 0.038 | −0.61 | 0.54 |
| morph | 3.37 | 0.18 | 18.96 | <.001 |
| human_face_2 | −0.055 | 0.48 | −0.12 | 0.91 |
| human_face_3 | −0.33 | 0.48 | −0.68 | 0.50 |
| color_change_condition_pulse | 0.016 | 0.050 | 0.32 | 0.75 |
| body_listening | −0.028 | 0.19 | −0.15 | 0.88 |
| noticing | −0.19 | 0.21 | −0.90 | 0.37 |
| attention_regulation | 0.29 | 0.21 | 1.40 | 0.16 |
| emotional_awareness | 0.10 | 0.17 | 0.60 | 0.55 |
| body_listening:color_change_condition_pulse | −0.063 | 0.082 | −0.77 | 0.44 |
| noticing:color_change_condition_pulse | 0.008 | 0.094 | 0.088 | 0.93 |
| attention_regulation:color_change_condition_pulse | 0.12 | 0.092 | 1.31 | 0.19 |
| emotional_awareness:color_change_condition_pulse | −0.003 | 0.073 | −0.040 | 0.97 |

**Table 4.**
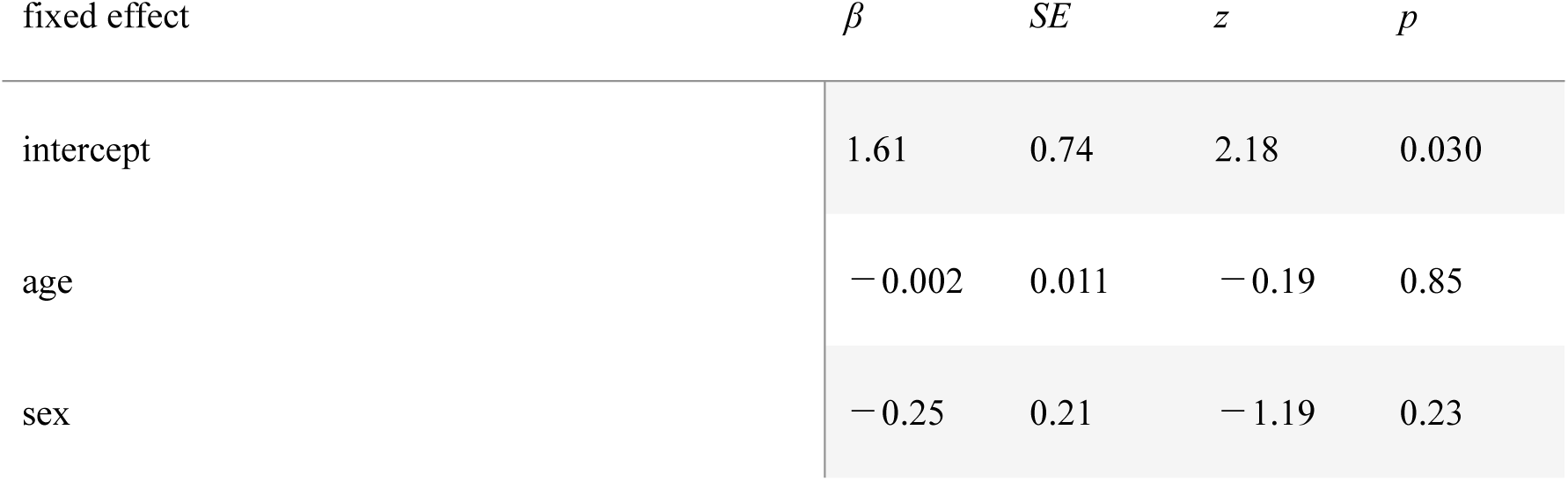
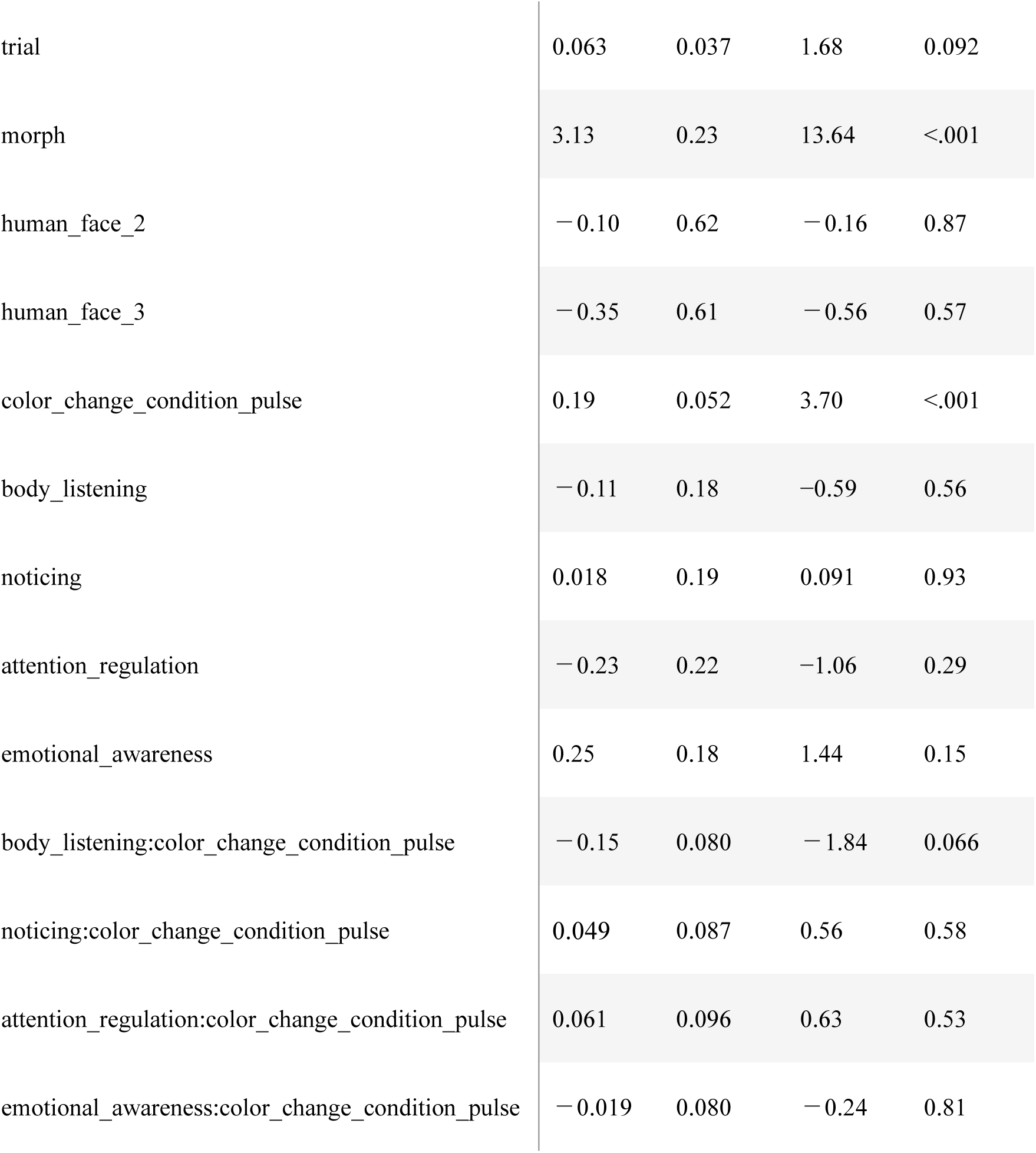
ERICA condition MAIA model.

| fixed effect | $\beta$ | $SE$ | $z$ | $p$ |
| --- | --- | --- | --- | --- |
| intercept | 1.61 | 0.74 | 2.18 | 0.030 |
| age | −0.002 | 0.011 | −0.19 | 0.85 |
| sex | −0.25 | 0.21 | −1.19 | 0.23 |
| trial | 0.063 | 0.037 | 1.68 | 0.092 |
| morph | 3.13 | 0.23 | 13.64 | <.001 |
| human_face_2 | −0.10 | 0.62 | −0.16 | 0.87 |
| human_face_3 | −0.35 | 0.61 | −0.56 | 0.57 |
| color_change_condition_pulse | 0.19 | 0.052 | 3.70 | <.001 |
| body_listening | −0.11 | 0.18 | −0.59 | 0.56 |
| noticing | 0.018 | 0.19 | 0.091 | 0.93 |
| attention_regulation | −0.23 | 0.22 | −1.06 | 0.29 |
| emotional_awareness | 0.25 | 0.18 | 1.44 | 0.15 |
| body_listening:color_change_condition_pulse | −0.15 | 0.080 | −1.84 | 0.066 |
| noticing:color_change_condition_pulse | 0.049 | 0.087 | 0.56 | 0.58 |
| attention_regulation:color_change_condition_pulse | 0.061 | 0.096 | 0.63 | 0.53 |
| emotional_awareness:color_change_condition_pulse | −0.019 | 0.080 | −0.24 | 0.81 |

### Examining the moderating effect of awareness of facial skin color changes

As an exploratory analysis, we examined whether explicit awareness of the facial skin color changes moderated the effect of color-change condition. The interaction between awareness and color-change condition was not significant for either the CommU condition (*β* = −0.09, *z* = −0.76, *p* = .45) or the ERICA condition (*β* = 0.05, *z* = 0.50, *p* = .61). Thus, the effect of pulse-wave-derived facial skin color changes on human-likeness judgments did not appear to depend on whether participants explicitly noticed the changes. Detailed results are presented in Tables 5 and 6.

**Table 5.** CommU condition color-change awareness model.

| fixed effect | $\beta$ | $SE$ | $z$ | $p$ |
| --- | --- | --- | --- | --- |
| intercept | 0.93 | 0.84 | 1.11 | 0.27 |
| age | -0.017 | 0.011 | -1.55 | 0.12 |
| sex | -0.33 | 0.19 | -1.71 | 0.087 |
| trial | -0.024 | 0.038 | -0.63 | 0.53 |
| morph | 3.37 | 0.18 | 19.06 | <.001 |
| human_face_2 | -0.079 | 0.48 | -0.17 | 0.87 |
| human_face_3 | -0.39 | 0.48 | -0.82 | 0.41 |
| color_change_condition_pulse | 0.17 | 0.22 | 0.81 | 0.42 |
| awareness | 0.19 | 0.28 | 0.68 | 0.50 |
| awareness:color_change_condition_pulse | -0.089 | 0.12 | -0.76 | 0.45 |

**Table 6.**
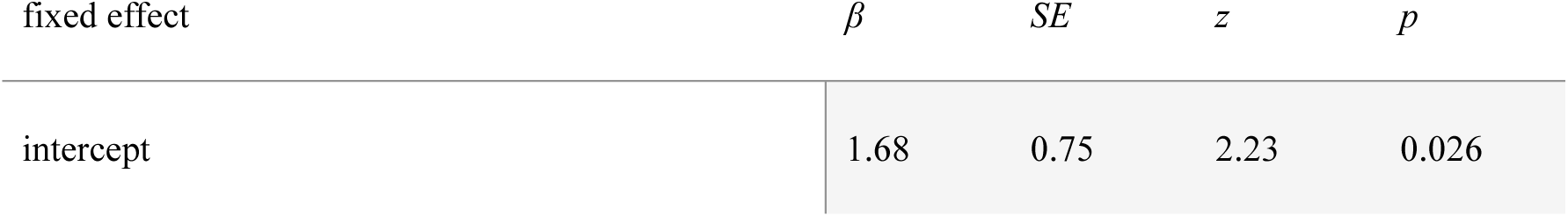
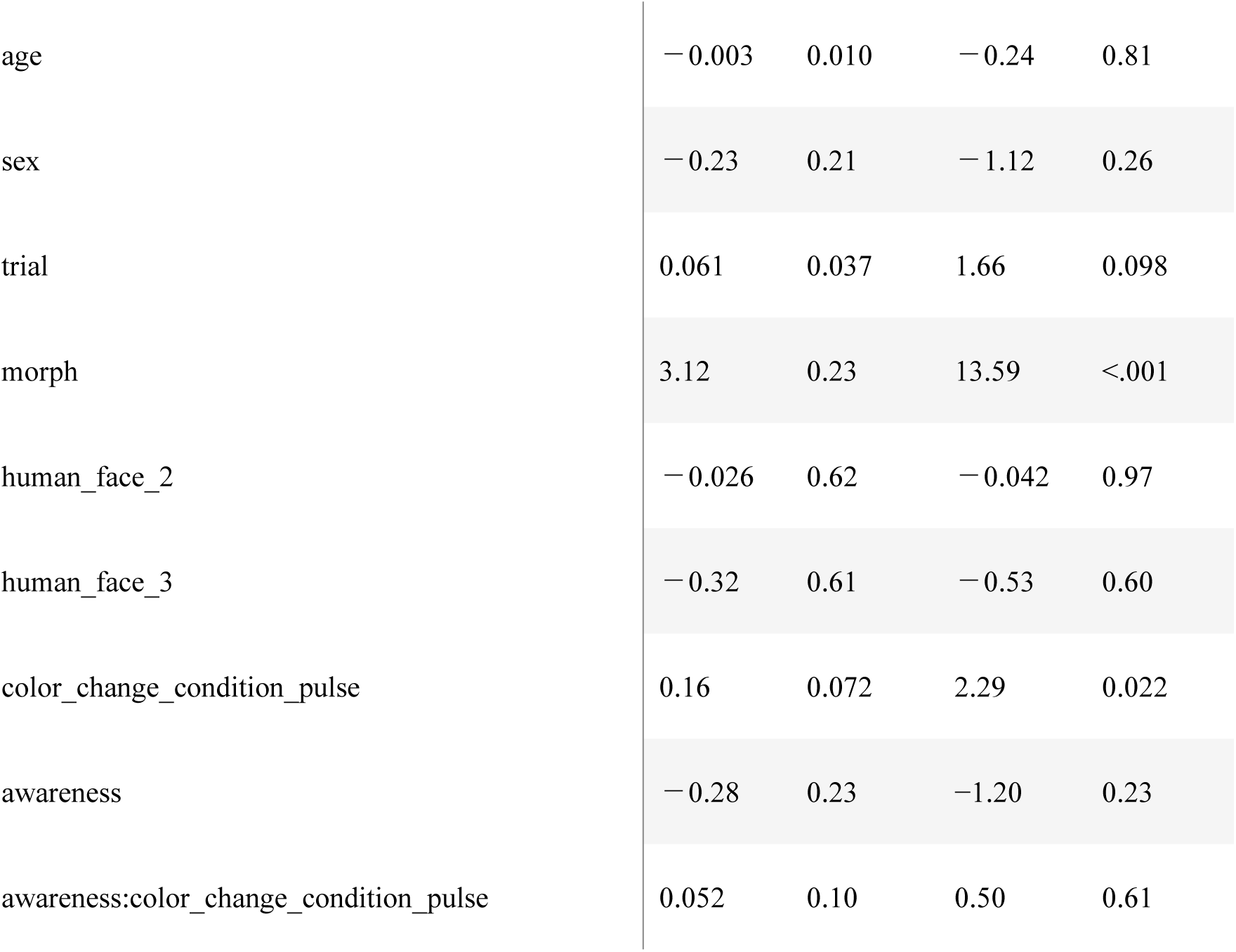
ERICA condition color-change awareness model.

Experiment 1 revealed that pulse-wave-derived skin color changes enhanced the perceived human likeness only for ERICA, a highly human-like agent. Additionally, the magnitude of this effect was not significantly moderated by interoceptive sensibility or awareness of the facial skin color changes. However, it remained unclear whether this effect was specific to genuine human pulse-wave signals or whether any rhythmic signal capable of inducing facial skin color changes would produce a similar effect.

### Experiment 2

The aim of Experiment 2 was to replicate the findings of Experiment 1 and to test whether human pulse-wave signals produced a stronger effect than non-biological rhythmic signals. To address this question, we introduced a sinusoidal condition in which facial skin color changed according to an artificially generated sinusoidal signal. Unless otherwise noted, the stimuli, procedure, and measures were identical to those used in Experiment 1.

## Methods

### Participants

Participants were recruited through CrowdWorks. A total of 212 participants completed the task. Of these, 67 participants were excluded because they had previously participated in Experiment 1. An additional 4 participants were excluded based on the morph level–human likeness judgment correlation criterion described in Experiment 1. The final sample therefore comprised 141 participants (57 men and 84 women; mean age = 41.18 years, SD = 11.29).

### Stimuli

Experiment 2 focused exclusively on ERICA. In addition to the pulse-wave and no-pulse conditions used in Experiment 1, a sinusoidal condition was introduced in which skin color changes were driven by a sinusoidal signal matched to the recorded pulse-wave signal in mean amplitude and number of peaks (Figure 2). Because little variation in judgment patterns across human face identities was observed in Experiment 1, a single human face was used for all participants.

**Figure 2.**
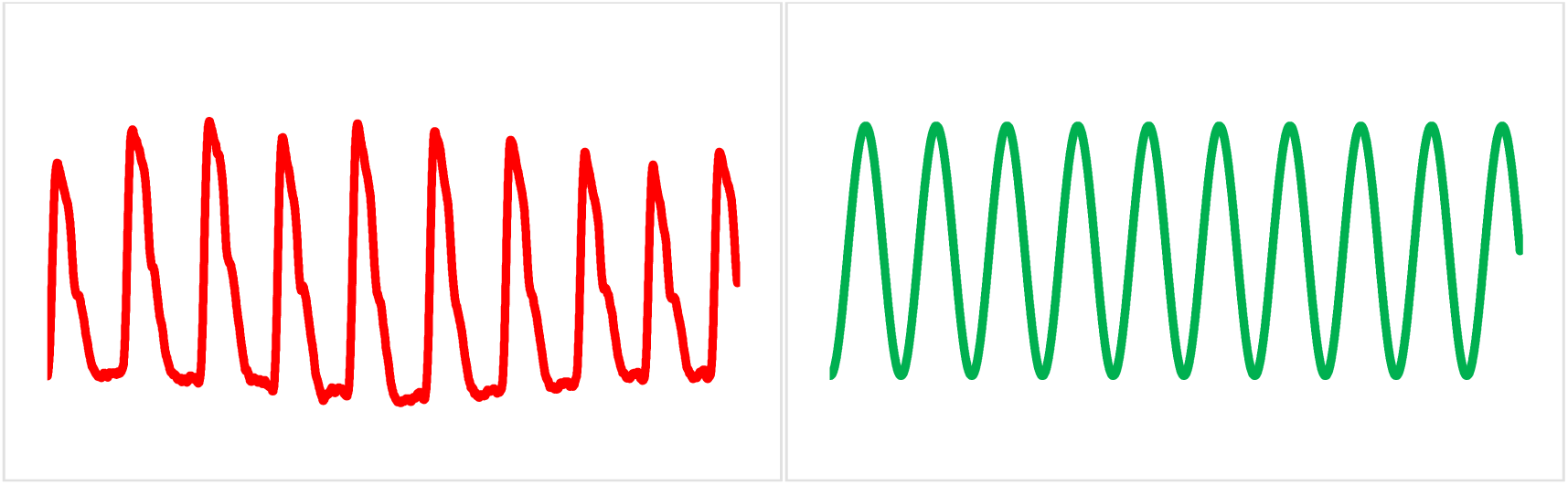
Waveforms of the signals used in Experiment 2. Left: pulse-wave signal used in the pulse-wave condition. Right: sinusoidal signal used in the sinusoidal condition.

### Procedure

The task followed the same procedure as in Experiment 1, except that the human-likeness judgment task included three color-change conditions: pulse-wave, sinusoidal, and no-pulse. In each block, participants viewed 21 video stimuli comprising 3 color-change conditions x 7 morph levels. Each stimulus lasted 8 s, after which participants judged whether the target appeared “human-like” or “robot-like”. The experiment consisted of four blocks, with stimulus order randomized within each block. Device restrictions, viewing instructions, and experimental platform were identical to those used in Experiment 1. Following the human-likeness judgment task, participants completed the MAIA-J and answered the same two post-task questions as in Experiment 1, concerning which aspects of the stimuli they had focused on when making their judgments and whether they had noticed any changes in facial skin color.

### Data Analysis

As in Experiment 1, we fitted generalized linear mixed-effects models (GLMMs) with a binomial distribution and logit link function to examine whether facial skin color changes enhanced perceived human likeness. We also examined whether interoceptive sensibility and explicit awareness of skin-color changes moderated the effect of color-change condition on human-likeness judgments. The model structures were identical to those used in Experiment 1, except that human-face identity was omitted because a single human face was used, and color-change condition consisted of three levels: pulse-wave, sinusoidal, and no-pulse. A likelihood ratio test (LRT) was used to assess whether adding color-change condition improved model fit. Pairwise comparisons among the three color-change conditions were conducted using estimated marginal means with Holm adjustment for multiple comparisons, implemented via the emmeans package in R (Lenth et al., 2026).

## Results

### Testing the Effect of Pulse-Wave-Derived Facial Skin Color Changes

Adding color-change condition significantly improved model fit relative to the baseline model (χ²(2) = 7.91, *p* = .02). In the main-effects model, morph level had a significant positive effect on human-likeness judgments (*β* = 3.25, *z* = 9.99, *p* < .001), indicating that stimuli were more likely to be judged as human as the proportion of the human face increased. More importantly, color-change condition significantly affected human-likeness judgments. Stimuli with pulse-wave-derived skin color changes were significantly more likely to be judged as human than stimuli without skin color changes (OR = 1.19, 95% CI [1.01, 1.42], *p* = .038). Similarly, stimuli with sinusoidal skin color changes were significantly more likely to be judged as human than stimuli without skin color changes (OR = 1.20, 95% CI [1.01, 1.42], *p* = .038). In contrast, no significant difference was observed between the pulse-wave and sinusoidal conditions (OR = 1.00, 95% CI [0.84, 1.19], *p* = .97).

These results suggest that the enhancement of perceived human likeness associated with pulse-wave-derived skin color changes may be attributable not to the specific temporal pattern of the pulse itself, but rather to the presence of periodic facial skin color changes more generally. Detailed results are presented in Table 7.

**Table 7.** ERICA condition main-effects model (Experiment 2)

| fixed effect | $\beta$ | $SE$ | $z$ | $p$ |
| --- | --- | --- | --- | --- |
| intercept | 0.49 | 0.85 | 0.59 | 0.56 |
| age | 0.007 | 0.013 | 0.54 | 0.59 |
| sex | -0.17 | 0.30 | -0.57 | 0.57 |
| trial | 0.098 | 0.053 | 1.86 | 0.063 |
| morph | 3.25 | 0.33 | 9.99 | <.001 |
| color_change_condition_pulse | 0.18 | 0.072 | 2.46 | 0.014 |
| color_change_condition_sin | 0.18 | 0.072 | 2.50 | 0.013 |

*Note.* The $p$ values are based on unadjusted Wald $z$ tests.

### Examining the moderating effect of interoceptive sensibility

We then tested whether individual differences in interoceptive sensibility moderated the effect of color-change condition on human-likeness judgments. Across all four subscales—Body Listening, Noticing, Attention Regulation, and Emotional Awareness—none of the interactions with either the pulse-wave or sinusoidal condition reached statistical significance. Thus, interoceptive sensibility did not significantly moderate the effects of either pulse-wave-derived or sinusoidal skin color changes on human-likeness judgments. Detailed results are presented in Table 8.

**Table 8.** ERICA condition MAIA model (Experiment 2)

| fixed effect | $\beta$ | $SE$ | $z$ | $p$ |
| --- | --- | --- | --- | --- |
| intercept | 0.52 | 0.84 | 0.61 | 0.54 |
| age | 0.007 | 0.013 | 0.56 | 0.58 |
| sex | −0.19 | 0.30 | −0.62 | 0.53 |
| trial | 0.099 | 0.053 | 1.87 | 0.062 |
| morph | 3.25 | 0.33 | 9.97 | <.001 |
| color_change_condition_pulse | 0.17 | 0.072 | 2.40 | 0.017 |
| color_change_condition_sin | 0.18 | 0.072 | 2.44 | 0.015 |
| body_listening | 0.095 | 0.23 | 0.41 | 0.68 |
| noticing | 0.53 | 0.27 | 1.96 | 0.050 |
| attention_regulation | − 0.42 | 0.29 | − 1.43 | 0.15 |
| emotional_awareness | − 0.35 | 0.21 | − 1.71 | 0.088 |
| body_listening: color_change_condition_pulse | − 0.048 | 0.11 | − 0.43 | 0.67 |
| body_listening: color_change_condition_sin | − 0.11 | 0.11 | − 1.01 | 0.31 |
| noticing: color_change_condition_pulse | − 0.13 | 0.13 | − 1.01 | 0.31 |
| noticing: color_change_condition_sin | − 0.13 | 0.13 | − 1.01 | 0.31 |
| attention_regulation:color_change_condition_pulse | 0.092 | 0.14 | 0.65 | 0.52 |
| attention_regulation:color_change_condition_sin | 0.11 | 0.14 | 0.78 | 0.44 |
| emotional_awareness:color_change_condition_pulse | 0.093 | 0.098 | 0.95 | 0.34 |
| emotional_awareness:color_change_condition_sin | 0.079 | 0.098 | 0.81 | 0.42 |

### Examining the moderating effect of awareness of facial skin color changes

As an exploratory analysis, we examined whether explicit awareness of the facial skin color changes moderated the effect of color-change condition on human-likeness judgments. Neither the interaction between awareness and the pulse-wave condition (*β* = 0.17, *z* = 1.09, *p* = .28) nor the interaction between awareness and the sinusoidal condition (*β* = 0.21, *z* = 1.31, *p* = .19) was statistically significant. Thus, there was no evidence that the effects of either pulse-wave-derived or sinusoidal skin color changes on human-likeness judgments depended on whether participants explicitly noticed the facial skin color changes. Detailed results are presented in Table 9. Table 9. ERICA condition color-change awareness model (Experiment 2)

**Table 9.** ERICA condition color-change awareness model (Experiment 2)

| fixed effect | $\beta$ | $SE$ | $z$ | $p$ |
| --- | --- | --- | --- | --- |
| intercept | 0.55 | 0.85 | 0.65 | 0.52 |
| age | 0.009 | 0.013 | 0.66 | 0.51 |
| sex | -0.17 | 0.30 | -0.58 | 0.56 |
| trial | 0.098 | 0.053 | 1.85 | 0.064 |
| morph | 3.25 | 0.33 | 10.00 | <.001 |
| color_change_condition_pulse | 0.13 | 0.086 | 1.47 | 0.14 |
| color_change_condition_sin | 0.12 | 0.086 | 1.38 | 0.17 |
| awareness | -0.40 | 0.34 | -1.18 | 0.24 |
| awareness:color_change_condition_pulse | 0.17 | 0.16 | 1.09 | 0.28 |
| awareness:color_change_condition_sin | 0.21 | 0.16 | 1.31 | 0.19 |

## General Discussion

The present study examined whether subtle facial skin color changes enhance the perceived human likeness of artificial agents and whether this effect depends on the type of agent, temporal characteristics of the signal or individual differences in interoceptive sensibility. In Experiment 1, pulse-wave-derived skin color changes enhanced human-likeness judgments for ERICA, a highly human-like agent, but not for CommU, a less human-like agent. In Experiment 2, both pulse-wave-derived and sinusoidal skin color changes increased human-likeness judgments relative to the no-pulse condition, whereas no difference was observed between the two dynamic conditions. Interoceptive sensibility and explicit awareness of the skin-color changes did not significantly moderate these effects.

We observed different effects of facial skin color changes on perceived human likeness between CommU and ERICA. This difference may be attributed to the relative salience of static morphological cues in human-likeness judgments. In the CommU condition, morphological changes in facial features such as the eyes and mouth were relatively pronounced across morph levels, which may have allowed participants to rely more heavily on static facial features when making their judgments. In contrast, morphological differences across morph levels were more subtle in the ERICA condition, potentially making judgments based on static information alone more difficult. Under these circumstances, dynamic cues such as facial skin color changes may have carried greater relative weight in judgments of human likeness. A complementary possibility is that the same dynamic cue was interpreted differently depending on the agent’s appearance. Previous research has suggested that the realism of an agent’s appearance can influence how dynamic cues are perceived and evaluated (Garau et al., 2003; Urgen et al., 2018). In ERICA, the facial color changes may have been more readily interpreted as cues to human physiological activity because of its highly human-like appearance. In contrast, the same changes may have been less diagnostic of human likeness when presented on CommU, whose appearance was more distinctly robotic.

The enhancement of perceived human likeness was not specific to actual pulse-wave signals, as a comparable effect was observed for sinusoidal skin color changes. These findings suggest that the fine-grained temporal structure specific to human pulse-wave signals may not be necessary for enhancing perceived human likeness. This pattern differs somewhat from McDuff and Nowara (2021), who reported stronger evaluations for pulse-wave-derived modulation than for sinusoidal modulation. This discrepancy may partly reflect differences in the evaluation task. Their task required relative judgments among simultaneously presented stimuli, whereas the present study assessed each stimulus independently using human/robot categorization. In addition, their pulse-wave and sinusoidal conditions differed not only in temporal structure but also in the manner of color modulation. By controlling the color-modulation procedure across signal types, the present study suggests that faithful reproduction of the human pulse waveform may not be necessary for enhancing human likeness. Importantly, however, the sinusoidal signals used in the present study were matched to the human pulse-wave signals in mean amplitude and number of peaks. Thus, the present results do not imply that any periodic color change would be sufficient; rather, changes occurring at a physiologically plausible magnitude and timescale may be important.

Contrary to our hypothesis, interoceptive sensibility did not significantly moderate the effect of facial skin color changes on human-likeness judgments. A possible reason is that the human/robot categorization task used in the present study primarily required judgments based on visual appearance. However, human likeness also encompasses social and psychological aspects, such as being perceived as an agent capable of experiencing emotions, having intentions, and communicating with others. The present task did not require participants to infer such internal states, which may have limited the extent to which individual differences in interoceptive sensibility influenced their judgments. In addition, the pulse-wave signals used in the present study were recorded at rest, and the stimulus videos were relatively short, lasting only 8 s. Thus, although the skin color changes may have conveyed cues to biological activity, they may not have provided sufficient information for observers to infer specific internal states such as arousal or emotion.

Rather than prompting explicit inferences about specific internal states, the skin color changes may have conveyed a more general impression of biological activity or aliveness. One possible mechanism underlying such an effect is animacy perception. Animacy can be perceived even from relatively simple visual cues when changes appear to originate from within an entity rather than from external physical causes (Tremoulet & Feldman, 2000; Scholl & Tremoulet, 2000). From this perspective, the periodic facial skin color changes used in the present study may have been perceived as autonomous changes arising from physiological activity within the agent, rather than as changes caused by external factors such as lighting or shadows. As a result, these changes may have enhanced the impression that the agent was biologically active, thereby influencing judgments of human likeness even without the inference of specific internal states such as emotions or intentions. Indeed, previous research has shown that not only object motion but also periodic temporal changes in luminance can enhance animacy perception (Inaba et al., 2016). Notably, animacy perception is considered to arise relatively automatically from visual stimuli, without necessarily requiring conscious inference or deliberate reasoning (Scholl & Tremoulet, 2000). This possibility is consistent with the present finding that explicit awareness of skin color changes did not significantly moderate their effect on human-likeness judgments. Thus, subtle periodic facial skin color changes may have served as cues to animacy even when participants did not consciously notice the changes. However, this interpretation remains tentative because the present study did not directly assess animacy perception.

Several limitations should be addressed in future research. First, the present study assessed only binary human/robot judgments. Future studies should examine whether facial skin color changes also influence other dimensions of agent perception, such as likability, friendliness, or attributed internal states. It will also be important to examine whether facial skin color changes affect other dimensions of perception in less human-like agents such as CommU, even when they do not influence human-likeness judgments. Second, the present study did not determine whether the observed effects were driven by periodicity itself or by temporal characteristics that were physiologically plausible. Future studies should therefore manipulate the frequency and amplitude of skin color changes across physiologically plausible and implausible ranges. Longer stimuli and signals reflecting changes in arousal and emotional state could also be used to test whether observers infer internal states from such physiological cues. Finally, examining whether human pulse-wave signals enhance animacy perception would help clarify whether perceived animacy mediates the effect of facial skin color changes on human-likeness judgments.

In summary, the present study showed that subtle facial skin color changes can enhance perceived human likeness, particularly for highly human-like artificial agents. This effect was observed not only for pulse-wave-derived signals but also for sinusoidal signals matched in mean amplitude and number of peaks, suggesting that faithful reproduction of the fine-grained temporal structure of human pulse waves may not be necessary. We also found that observers’ interoceptive sensibility did not moderate this effect. This may reflect the limited information available for inferring specific internal states from brief skin color changes based on resting pulse-wave signals. Together, these findings highlight the potential importance of subtle physiological-like facial dynamics in shaping perceptions of human likeness in artificial agents.

